# Fitting functional responses: Direct parameter estimation by simulating differential equations

**DOI:** 10.1101/201632

**Authors:** Benjamin Rosenbaum, Björn C. Rall

**Author notes:** Corresponding authors &.

## Abstract

1. The feeding functional response is one of the most widespread mathematical frameworks in Ecology, Marine Biology, Freshwater Biology, Microbiology and related scientific fields describing the resource-dependent uptake of a consumer. Since the exact knowledge of its parameters is crucial in order to predict, for example, the efficiency of biocontrol agents, population dynamics, food web structure and subsequently biodiversity, a trustful parameter estimation is of utmost importance for scientists using this framework. Classical approaches for estimating functional response parameters lack flexibility and can often only serve as approximation for a correct parameter estimation. Moreover, they do not allow to incorporate side effects such as resource growth or background mortality. Both call for a new method to be established solving these problems.
2. Here, we combined ordinary differential equation models (ODE models), that were numerically solved using computer simulations, with an iterative maximum likelihood fitting approach. We compared our method to classical approaches of fitting functional responses, using data both with and without additional resource growth and mortality.
3. We found that for classical functional response models, like the often used type II and type III functional response, the established fitting methods are reliable. However, using more complex and flexible functional responses, our new established method outperforms the traditional methods. Additionally, only our method allows to analyze experiments correctly when resources experience growth or background mortality.
4. Our method will enable researchers from different scientific fields that are measuring functional responses to estimate parameters correctly. These estimates will enable community ecologists to parameterize their models more precisely, allowing for a deeper understanding of complex ecological systems, and will increase the quality of ecological prediction models.

## Introduction

Understanding the interactions between organisms is crucial to understand patterns of stability and biodiversity of whole communities (McCann, 2000). Theory suggests that especially antagonistic interactions, such as feeding interactions, may have negative effects on stability (May, 1972) and subsequently on biodiversity (e.g. Rall, Guill & Brose, 2008). Moreover, a good knowledge on interactions may allow ecologists to create realistic predictions of empirical communities ranging from microcosms to whole lakes (Boit *et al.,* 2012; Schneider, Scheu & Brose, 2012; Fussmann *et al.*, 2014). It is therefore of utmost importance to have reliable methods for quantifying interaction strength amongst organisms, especially consumer-resource pairs such as predators and their prey.

### The functional response models

The functional response (Solomon, 1949; Holling, 1959b) is one of the oldest and most commonly-used mathematical frameworks to describe and estimate the feeding interaction between a consumer and a resource (see Jeschke, Kopp & Tollrian (2002) for an overview). It describes the resource density-dependent feeding rate of a consumer on its resource (Holling, 1959b). Despite the plethora of possibilities, the most important functional response models used today are the type II and the type III functional responses (Skalski & Gilliam, 2001; Jeschke, Kopp & Tollrian, 2004). The type II response is a hyperbolic satiating curve (Fig. 1, blue line) where the *per capita* feeding rate, *F*, depends on the number of prey in the environment, *N*:

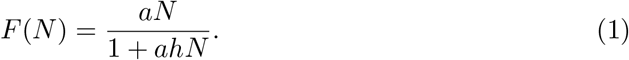

**Figure 1:**
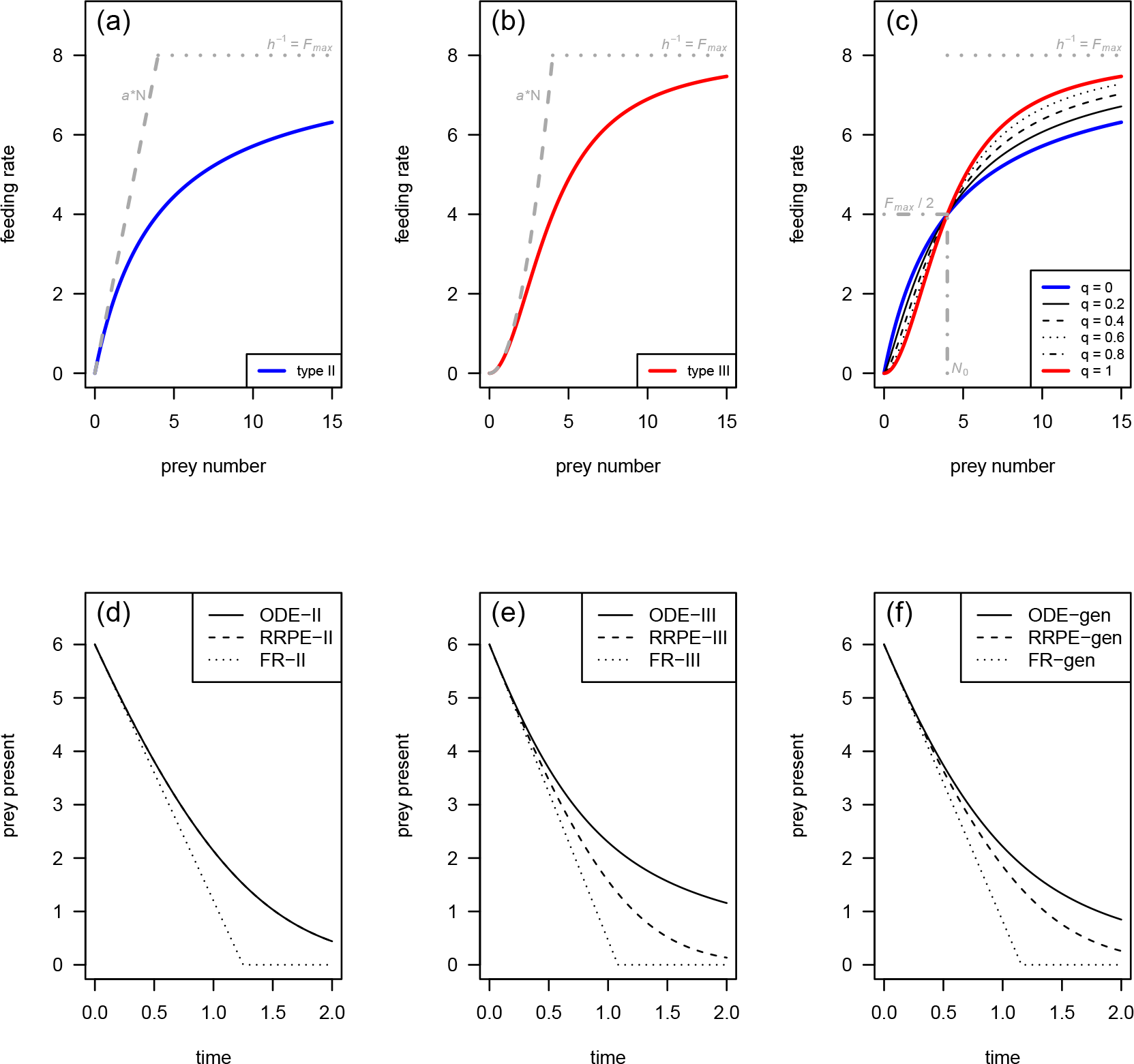
Top: Feeding rate *F*(*N*) for three different functional response models, Holling’s type II (a), Holling’s type III (b), and the generalized response (c). Bottom: Predicted number of prey present *N*(*t*) = *N*_0_ – *N*_*e*_(*t*) over time *t* for the type II (d), type III (e) and a generalized (*q* = 0.5) functional response (f), while ignoring prey depletion (dotted), using the RRPE (dashed) and our new numerical solution of the ODE (solid).

Here, *a* is the instantaneous rate of discovery (Holling, 1959b), commonly known and subsequently used here as attack rate (note that in dependence of the scientific field you may also know it as capture rate (e.g. Kalinkat *et al.*, 2011), maximum clearance rate (e.g. Hansen, Bjornsen & Hansen, 1997), maximum per capita interaction strength (e.g. McCann, Hastings & Huxel, 1998), or others); and *h* is the handling time (Holling, 1959a). See Jeschke, Kopp & Tollrian (2002, 2004) for a comprehensive introduction and discussion of the ecological meaning of these parameters. The attack rate, *a,* controls mainly the initial increase of feeding at low densities (Fig. 1a, gray dashed line) whereas the handling time, *h*, controls mainly the feeding at high densities where the feeding curve satiates (Fig. 1a, gray dotted line). The inverse of the handling time is often referred to by the maximum feeding rate, *F*_max_, but also by the maximum ingestion rate (e.g. Hansen, Bjornsen & Hansen, 1997), the handling rate (e.g. Englund *et al.*, 2011), or similar terms.

If the attack rate is not a constant but depends linearly on the resource density (*a* = *bN*), the functional response becomes a sigmoid curve (e.g. Juliano, 2001, see Fig. 1b, red line) that can be described by:

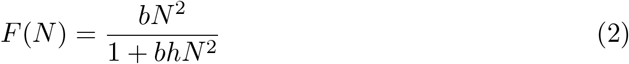

where *b* is the attack coefficient (please see Juliano (2001) for more possibilities to model a type III functional response). As for the type II functional response, the attack rate, *a*, controls the feeding rate at low prey densities (Fig. 1b, gray dashed line) and the handling time, *h*, controls the feeding rate at high prey densities (Fig. 1b, gray dotted line).

Real (1977, 1979) incorporated a possibility to gradually shift between the above mentioned fixed two types of the functional responses by using the enzyme kinetics model by Barcroft & Hill (1910). In Real’s adaptation, the attack rate depends with an power law on resource density (*a* = *bN*^*q*^), leading to functional response model that can be written as:

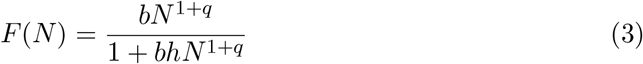

where *q* is a exponent influencing the shape of the functional response from a hyperbolic type II functional response (*q* = 0) to a strict type III functional response (*q* = 1) and also beyond these borders (Fig. 1c). Note that the original formulation of this functional response is 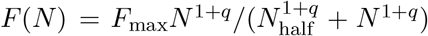, were *F*_max_ is the already above mentioned maximum feeding rate (*h* = 1/*F*_max_), and *N*_half_ is the half saturation density - the prey density at which the predator consumes half the amount of its maximum possible feeding rate 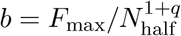; Fig. 1c, gray dotted-dashed line, please see the supplement (Sec. 1) for the equivalence of both parametrisations and Real (1977)). Moreover, the attack exponent, *q*, is often not used directly but in its transferable version, the Hill exponent, *H* (*H* = *q* + 1). This kind of functional response is also often referred to as generalized functional response (e.g. Kalinkat *et al.*, 2013; Barrios-O’Neill *et al.*, 2016). Using this elegant generalized functional response model revealed that increasing the attack exponent, *q*, just slightly introduces stability of simple consumer-resource population models up to whole food webs, and thereby increases biodiversity (Williams & Martinez, 2004; Rall, Guill & Brose, 2008; Uszko *et al.*, 2015). Inspired by these theoretical findings, researchers aimed to investigate which traits (e.g. body mass Vucic-Pestic *et al.*, 2010; Kalinkat *et al.*, 2013; Barrios-O’Neill *et al.*, 2016) or environmental drivers (e.g. temperature (Uszko *et al.*, 2017) or habitat structure (Barrios-O’Neill *et al.*, 2016; Li *et al.*, 2017)) change the attack exponent, *q*, from being zero (a destabilizing type II functional response) to higher values (a stabilizing generalized functional response).

### The problem of prey depletion - and its possible solutions

The above mentioned functional response models (eqns. 1, 2, 3) describe how feeding rates depend on the number of prey in the environment (Fig. 1a-c). This means that the independent variable (displayed on the x-axis) must be a constant prey density. In experiments, however, it is hardly possible to keep the prey density constant over time. Most feeding trials with the aim of estimating a functional response are principally designed as follows: (1) a selected number of resource items is introduced into an experimental arena; (2) one or more consumers are added; (3) the experiment is run over a defined time range; (4) the remaining prey items are counted; (5) the number of prey remaining is subtracted from the initial number of prey, yielding the number of prey consumed over time. This has two implications: (1) the measured variable is not a rate, but the cumulative number of eaten prey items; and (2) the independent variable - the prey density - decreases over time. This phenomenon is commonly known as prey depletion (Rogers, 1972).

Describing the change of the population density over time is classically done by using ordinary differential equations (ODE); this approach is applied to model a wide range of ecological systems, from single populations up to whole multi-trophic communities (e.g. Verhulst, 1838; Rosenzweig & Mac Arthur, 1963; Rall, Guill & Brose, 2008; Delmas *et al.*, 2016). To correctly describe the prey depletion during the course of a functional response experiment, we can set up such an equation using the generalized Holling functional response (eqn. 3):

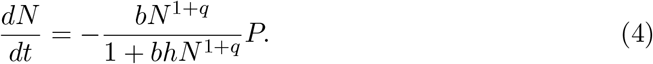

This equation describes the change of the prey density, *dN*, over time *dt*, which is proportional to the feeding rate, i.e. the generalized functional response (eqn. 3), multiplied the predator density, *P*. ODE models must be integrated over time and density to yield the density at a desired point in time - here, the prey density at the end of the experiment, *N*_end_ (Fig. 1d-f, black solid lines). This integration process can be done analytically (i.e. using mathematical rules to derive a new function having time as an independent variable and density as a dependent variable); or, if the ODE is too complex to be solved analytically, it can be solved numerically using computer algorithms that approximate the real solution, with an accuracy that it is not distinguishable from that of an analytical solution (i.e. computer simulations). After integration the number of prey eaten over time, *N*_*e*_, is calculated by simply subtracting the remaining prey density at the end of the simulation (as for experimental data), *N*_*end*_, from the starting density, *N*_0_ (*N*_*e*_ = *N*_0_ – *N*_*end*_). As mentioned above, this generalized equation (4) becomes a type II functional response if the attack exponent is zero (*q* = 0, Fig. 1d), and a type III functional response if the attack exponent becomes unity (*q* = 1, Fig. 1e). In the following, we present established methods, which use the functional response to predict the number of eaten prey *N*_*e*_.

The most simple approximation that could be made to describe the depletion of prey over the course of the experiment is to multiply the feeding rate, *F*, by the number of predators (more generally: consumers) and the time of the experiment, *T*, yielding the number of prey eaten at the end of the experiment, *N*_e_:

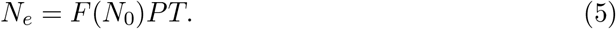

Here, the calculation of the feeding rate, *F*, can replaced by any of the above mentioned functional response models (eqns. 1–3) under the constraint that the density, *N*, is replaced by the starting number of prey items, *N*_0_. This method is a linearization of the non-linear process that is precisely described by the ODE (eqn. 4) and yields a linear decrease of alive prey in the experiment over time (Fig. 1, d–f, dotted lines).

Instead of linearizing the ODE (eqn. 4), Royama (1971) and Rogers (1972) presented an analytical solution of the ODE incorporating a type II functional response (*q* = 0), commonly known as Rogers Random Predator Equation (RRPE, for the analytical derivation see supplement, Sec. 2):

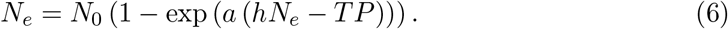

The number of prey consumed at the end of the experiment, *N*_*e*_, depend here on the regular functional response parameters attack rate, *a*, and handling time, *h*, as well as the total time of the experiment, *T*, and the number of predators, *P*. This equation is commonly used to estimate the parameter values for a type II functional response when prey depletion occurs, and we will refer to it as Rogers Random Predator Equation II (RRPE-II).

With the unknown number eaten, *N*_*e*_, appearing on both sides of this implicit equation (RRPE-II), the problem arises that a simple non-linear fitting algorithm is not applicable. The most frequently used solution to this problem is to use an iterative Newton’s method (Juliano & Williams, 1987; Juliano, 2001). However, the state-of-the-art solution is to use the LambertW function instead, which allows an explicit solution of the implicit RRPE-II (Bolker, 2008):

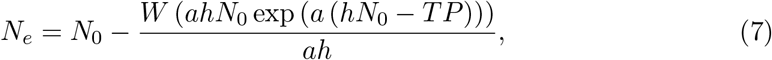

see also Fig. 1d, dashed line. This approach has already been incorporated into a R-package for easy application (Pritchard, 2016; Pritchard *et al.*, 2017).

Juliano (2001) suggested that for fitting type III functional responses with prey depletion, it is possible to simply include the resource density dependency of the attack rate and replace the time-dependent prey number, *N*, by the constant value of the prey number at the start of the experiment, *N*_0_ (*a* = *bN* ≈ *bN*_0_), into the RRPE-II yielding:

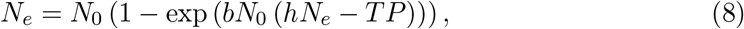

where the parameters are as described above: the number eaten, *N*_*e*_, the attack constant, *b*, the handling time, *h*, the total time, *T*, and the number of predators *P*. We will subsequently refer to this approach as Rogers Random Predator Equation III (RRPE-III).

Just as the RRPE-II, this implicit equation can be solved iteratively by Newton’s method for *N*_*e*_ or by using the LambertW function:

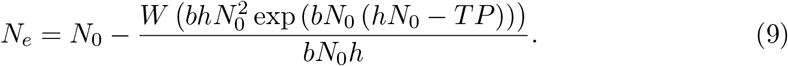

Regardless of whether this equation is solved implicitly or explicitly, due to the approximation *a* = *bN* ≈ *bN*_0_ the predicted number of eaten prey *N*_*e*_ is an approximation to the type III ODE and not an analytically correct solution (Fig. 1e, dashed line).

As an alternative to the type III functional response, Hassell, Lawton & Beddington (1977) presented a sigmoid-shaped feeding curve, taking prey depletion into account. Since the classical type III response is a special case of their wider class of curves (see supplement, Sec. 3), we derived a simplified version of their original equation describing how prey will be depleted over time following a type III functional response:

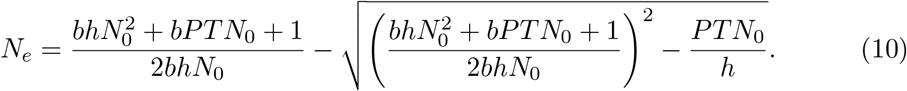

This equation represents an analytically correct solution of an ODE with a type III functional response (for the derivation see supplement, Sec. 3). This function is more complex as the RRPE-III, but it contains exactly the same parameters: the number eaten, *N*_*e*_, the attack constant, *b*, the handling time, *h*, the total time, *T*, and the number of predators *P*. But it has the advantage that the dependent variable, *N*_*e*_, is only on the left side of the equation, allowing standard fitting algorithms to be used.

While the above models can be used to fit either type II or type III functional responses, the RRPE can also be used to fit the generalized functional response (eqn. 3) to data with prey depletion. The power law dependency of the attack rate, 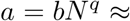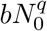 was incorporated into the RRPE-II (e.g. Vucic-Pestic *et al.*, 2010; Kalinkat *et al.*, 2013; Barrios-O’Neill *et al.*, 2016), yielding:

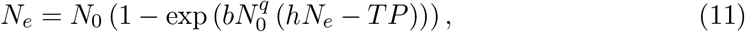

where the parameters are as above: the number eaten, *N*_*e*_, the attack constant, *b*, the attack exponent, *q*, the handling time, *h*, the total time, *T*, and the number of predators *P*. We will subsequently refer to this approach as generalized Roger’s Random Predator Equation (RRPE-gen).

Again, this implicit problem can be solved iteratively by Newton’s method or explicitly by using the LambertW function

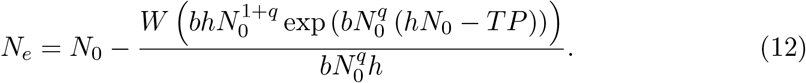

Analogous to the RRPE-III, these solutions of the RRPE-gen are an approximation to the ODE with a generalized FR (Fig. 1f, dashed line).

As an alternative, Uszko *et al.* (2015, 2017) generalized a method developed by Frost (1972) in which (1) an approximation of an average resource density, 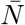, is calculated 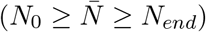 - taking prey depletion into account - and (2) the functional response parameters are estimated using (for further information see supplement, Sec. 4):

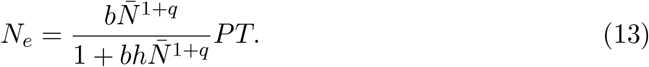

### Possible drawbacks of the different methods

Using the most simple approach where the feeding rate is simply multiplied by predator density and time of the experiment (eqn. 5), prey depletion during the time of the experiment is completely neglected. Because this feeding rate is higher than the actual feeding rate that is based on a reduced number of prey later in the experiment (*N*_0_ > *N* causes *F*(*N*_0_) > *F*(*N*), cf. Fig. 1d-f, compare the dotted lines to the solid line created by the ODE representing the natural depletion of prey), attack rates, *a*, will be underestimated, regardless whether a type II, a type III or a generalized functional response is used. As the feeding rate as well as the realized feeding over the course of an experiment (i.e. the total amount of consumed prey) approaches a satiation level, the estimate of the maximum feeding rate *F*_max_ (or the handling time 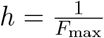) should only be marginally affected.

The RRPE-II is the exact analytical solution of an ODE that describes the depletion of prey over time with a type II functional response. The resulting depletion of prey over time is consequently identical if using the RRPE-II or the numerical simulation of an ODE (Fig. 1d, dashed vs. solid line). This method therefore delivers reliable parameter estimates if the data can be described by a type II functional response.

As the RRPE-II is the exact solution for an ODE with a type II functional response, there can be negative consequences when it is used to fit non-type II functional responses, such as the type III and the generalized functional response. The attack rate, *a* = *bN*^*q*^, in this case is itself prey number-dependent. Approximating *N* by the larger *N*_0_ creates a larger attack rate and therefore a stronger decrease of prey than given in a simulated ODE (Fig. 1e–f, dashed vs. solid line). A stronger decrease of prey should lead to an underestimation of the attack rates (i.e. attack coefficient *b*), whereas the handling time should be estimated correctly as both methods produce a similar linear decrease of prey density at high prey densities. An increasing attack exponent, *q*, decreases the number of prey consumed at low prey numbers, as prey depletion is predicted to be too high by using the Rogers Random Predator equation with a type III or generalized functional response. We assume that the attack exponent, *q*, would be overestimated to counteract this effect.

We were not able to create strong expectations for the method by Frost (1972) and Uszko *et al.* (2015) since this 2-step process can not be visualized as descriptively as the previous methods. But as the original method is based on a type I functional response, we assume also deviations from the correct solution in all cases.

Using our new analytical solution for a type III functional response (eqn. 10) will result in a correct estimation of functional response parameters. But more importantly, our method to fit numerical simulations of ODEs allows to include different functional response models and will always return correct parameter estimations.

### The problem of non-consumer mediated prey growth and mortality

Another problem that often arises in experiments is that background mortality or growth of the prey could occur. For example, the assumption of constant prey numbers of the experiment could be violated not only because of prey removal by predators, but also because of prey mortality, or population growth, what is often the case if using microbial or algal prey (e.g. Fussmann *et al.*, 2017; Uszko *et al.*, 2017). Standard ways to deal with this are to exclude any data that is biased with mortality or growth, leading to a substantial loss of data, or to ignore the problem if it is not too pronounced, accepting that the resulting bias in parameters is likely to be small (e.g. Vucic-Pestic *et al.*, 2010; Uszko *et al.*, 2017). However, it is also possible to minimize bias without excluding data, by fitting the parameters using simulations of ODEs (see e.g. Rall & Latz, 2016). Additional mortality can be accounted for by adding a mortality rate, *m*, to the ODE (eqn. 4). This a standard way to model mortality in population and food web modeling (e.g. Fussmann *et al.*, 2014; Kalinkat *et al.*, 2013; Williams & Martinez, 2004; Rosenzweig & Mac Arthur, 1963; Rall, Guill & Brose, 2008)

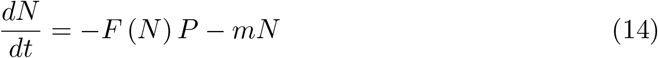

for arbitrary functional response models *F*(*N*).

The growth of prey is often described by simple phenomenological growth models
such as the logistic growth (Verhulst, 1838) or by Gompertz growth model (Gompertz, 1825; Paine *et al.*, 2012). The logistic growth can be added to the ODE model by simply adding the growth term as the mortality above. In the case of the logistic growth the ODE becomes:

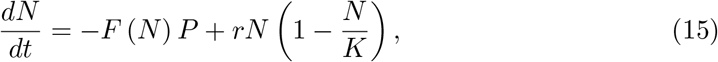

where *K* is the carrying capacity, the number of prey a system can sustain, and *r*, the intrinsic growth rate that controls the prey growth at low prey densities but also determines the mortality of prey if the prey density is above the carrying capacity. The rest of the parameters are as above in the case of a simple functional response ODE.

In the cases of both prey mortality and combined prey growth and mortality, we expect biased functional response parameters if the estimation is not corrected for these additional effects. Assuming experiments where prey mortality occurs (eqn. 14, e.g. using insects, spiders or crustaceans) this could lead to an overestimation of the attack rate or the attack coefficient, an underestimation of the attack exponent (the lower the exponent, the higher the feeding at low prey numbers), and an underestimation of the handling time (the lower the handling time, the higher the maximum consumed number of prey) to counteract the additional effect of mortality. In the case where the prey is growing and dying according to the above mentioned growth model (eqn. 15), there would likely be an underestimation of attack rate (which describes feeding at low prey numbers) or attack coefficient and potentially an overestimation of the attack exponent to counteract the additional growth of prey at low numbers. At high prey numbers exceeding the carrying capacity, background prey mortality occurs and there would likely be an underestimation of handling times (which describes feeding at high prey numbers) to counteract this effect. Using the extended ODE models and fitting them numerically to the data should overcome the problem. We further expect that using additional control data (no predators present) will improve the accuracy of the parameter estimation, because effects of natural growth (or death) and predation can be disentangled.

Here, we present a new framework of how to fit functional response models to data. We combine numerical simulations of ODE models with an iterative maximum likelihood estimator using **`R`** (Rall & Latz (2016), but see DeLong, Hanley & Vasseur (2014) for alternatives)This allows us to fit any functional response model to data more flexible and precisely than traditional methods. Our approach produces similar results if using the special case of a strict type II functional response as Juliano’s and Bolker’s method that use Rogers Random Predator Equation. Also, we expect that in the special case of a "strict" type III functional response, the parameter estimation by the analytical solution and numerical simulation methods should deliver similar results. In the case of the generalized functional response, which is widely used in current ecological studies (Vucic-Pestic *et al.*, 2010; Barrios-O’Neill *et al.*, 2016; Kalinkat *et al.*, 2013; Uszko *et al.*, 2015; Pawar, Dell & Savage, 2012; Li *et al.*, 2017) we expect that our method outperforms traditional methods dramatically. Moreover, fitting numerical simulations of ODE models also allows to cope with side effects like prey growth and mortality that otherwise may bias or even prevent the estimation of the functional response parameters.

## Materials and Methods

We simulated 1000 datasets based on ODE models (eqn. 4) for the type II, the type III and the generalized functional response each. For numerical simulation, we used the deSolve package (Soetaert, Petzoldt & Setzer, 2010). We drew attack coefficients, handling times and, in the generalized case, attack coefficients from random distributions (see supplement, Sec. 5). Each dataset contained in total 960 observations of eaten prey *N*_*e*_ from initial prey numbers *N*_0_ ranging from 1 to 48 individuals. We further simulated 1000 datasets for the generalized functional response including prey mortality (eqn. 14) and also 1000 datasets for the generalized functional response including prey growth (and natural prey death, if the prey number exceeds the carrying capacity, eqn. 15). Again, see the supplement (Sec. 5) for details on data simulation.

We used an iterative maximum likelihood method (bbmle package, function `mle2()`, Bolker (2008); Bolker & R Development Core Team (2016)) to fit the models to simulated data and also experimental datasets (see below). Unless stated otherwise, we assumed binomially distributed numbers of eaten prey. This means that for each observation 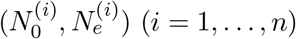, a prediction for the number of eaten prey 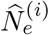 was calculated based on the respective model and its parameters. The observation 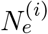 was then modeled as being binomially distributed with 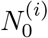 trials and a success probability of 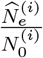 to compute the likelihood. See the manual (Sec. 6) for more information on the construction of likelihood functions and the fitting routines.

The growth model requires a different statistical approach. Through growth, the number of prey present at the end of the experiment *N*_*end*_ can exceed the number of initial prey *N*_0_. Negative numbers of eaten or dead prey *N*_*e*_ = *N*_0_ – *N*_*end*_ occur and make the binomial distribution unsuitable for this model. Instead, we use *N*_*end*_, which is always non-negative. For each observation 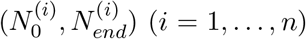 the prediction of the number of prey present 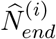 at the end of the experiment was calculated by numerical simulation of the ODE. The observation 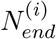 was modeled as being lognormally distributed with location parameter log 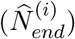 and scale parameter σ. This scale parameter is automatically estimated in the iterative maximum likelihood routine, too (Crawley, 2012, Chap. 7.3.3). The lognormal distribution takes into account that residuals generally have a larger variance for large prey abundances. See the manual (Sec. 6.6) for a detailed description of this likelihood function.

For the data without prey growth or mortality, we fitted all corresponding feeding models and compared the original simulation parameters to the estimates. For simulated data including background prey mortality, we tested the performance of fitting the ODE model neglecting mortality (eqn. 4) and the ODE model including mortality (eqn. 14). We also used the mortality model to fit simulated data which used half of the 960 observations as control data, i.e. no predators are present and prey death occurs by natural mortality only. For data including prey growth and mortality, we also tested the performance of fitting the ODE model neglecting growth (eqn. 4) and the ODE model including growth (eqn. 15). We also used the growth model to fit simulated data which used half of the 960 observations as control data, i.e. no death by predation occurs. We evaluated the performance all models by comparing the log_10_-ratios of estimated and true parameters (Berlow *et al.*, 1999; Rall & Latz, 2016).

We fitted our new ODE-based models (eqn. 4) to six experimental datasets from the literature and compared the results to the studies’ original methods, which are as follows. Sentis, Morisson & Boukal (2015) used the RRPE-III to fit dataset D1, while the RRPE-gen was fitted to dataset D2 in Vucic-Pestic *et al.* (2010). Uszko *et al.* (2015) used Frost’s method with a generalized FR for dataset D3. Note that this dataset consists of prey densities instead of prey individuals, which requires a lognormal distribution (for details see the manual, Sec. 6.5). Barrios-O’Neill *et al.* (2015) fitted the RRPE-gen to experimental data featuring a low (D4), medium (D5) or high (D6) spatial refuge for prey, which result in lower, medium and higher attack exponents.

The additional dataset D7 (L. Archer, 2017, personal communication) includes natural prey mortality and it also contains control data without any predators present. As with simulated data (see above), we compared the performance of the generalized ODE neglecting mortality (eqn. 4) to the models including mortality (eqn. 14) without and with using control data.

In a last example, we use dataset D8 (Fussmann, 2017) which features natural prey growth and death. The example also includes control data without predators present. Similar to the simulated data, we fit the generalized ODE neglecting growth (eqn. 4) and the model including a growth term (eqn. 15) with and without using control data.

## Results

### Feeding models

The results of fitting type II, type III and generalized functional response feeding models to simulated data are depicted in Fig. 2. For the data simulated with a **type II** response, our new ODE model (eqn. 4), and also the old models by Bolker (eqn. 7) and Juliano (eqn. 6) represent analytically correct feeding models. They provide an unbiased estimation of attack rates and handling times, i.e. the mean error over 1000 fits equals zero. Fitting the functional response term directly to the data (eqn. 5) leads to a heavily biased estimation, because the method does not correct for depleted prey. As expected, attack rates are systematically underestimated. Handling times are underestimated, too. When using Frost’s method for correction (eqn. 13), these errors are reduced, but the estimates are still biased.

**Figure 2:**
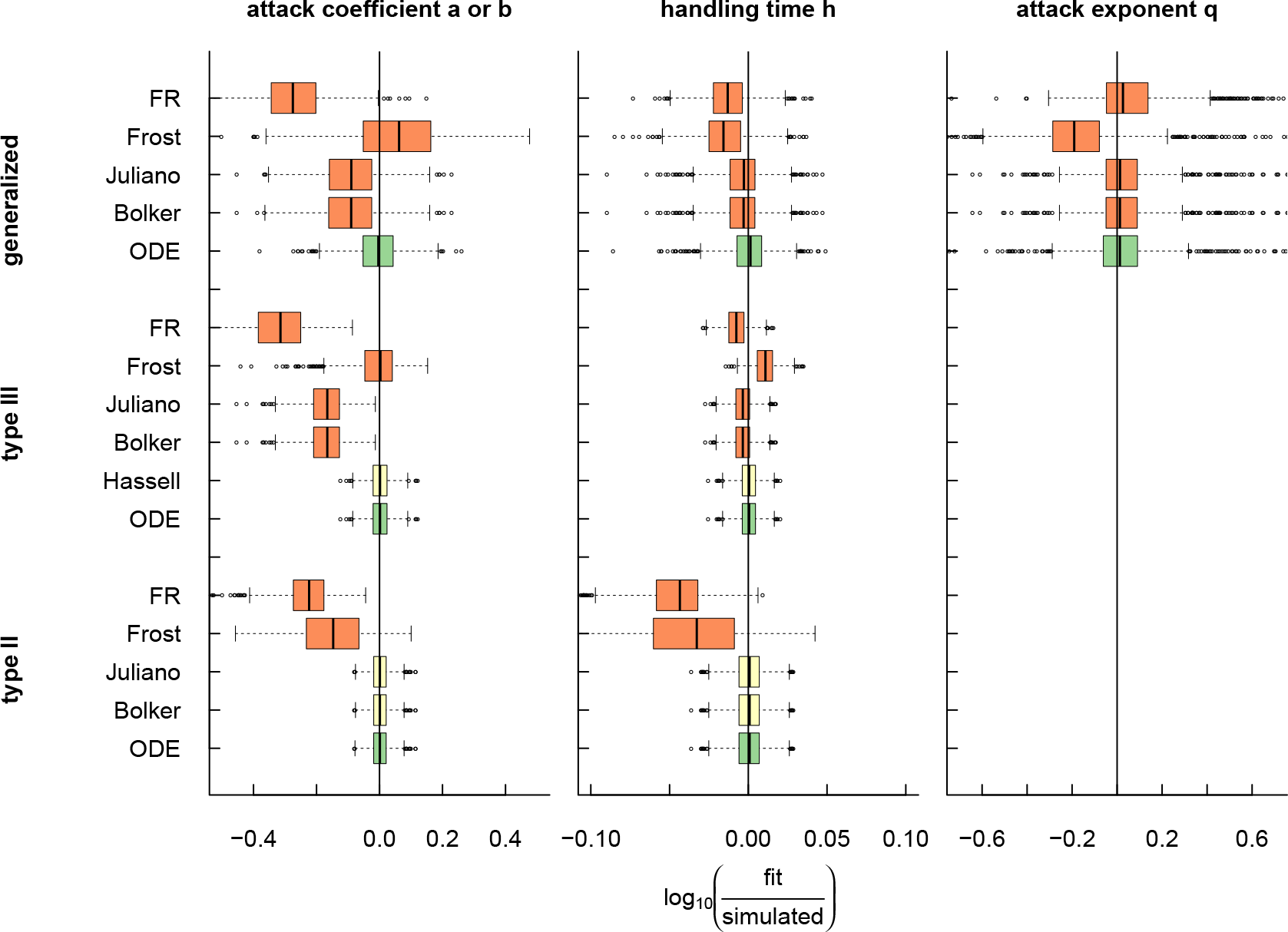
Evaluation of feeding models for the type II, type III and generalized functional response. The boxplots show the error distributions of fitted parameters 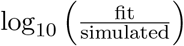 in 1000 simulated datasets. Green and yellow indicate analytically correct feeding models, orange indicates approximations.

In the case of a **type III** response, our new ODE model (eqn. 4) and also the model based on Hassell’s method (eqn. 10) provide analytically correct solutions for the number of eaten prey. Both return unbiased estimates for attack rates and handling times. For the method of Bolker (eqn. 9) and Juliano (eqn. 8), the approximation of *a* = *bN* by *bN*_0_ causes a severe underestimation of attack coefficients. Also, the handling time estimates are biased. Again, a direct fit of the functional response to the data (eqn. 5) leads to a heavy underestimation of attack coefficients as well as biased handling times. While Frost’s correction for depleted prey (eqn. 13) features an overall unbiased estimation of attack coefficients, the spread of errors is comparably large and handling times estimates are still biased.

For a **generalized** model with flexible attack exponent, our new ODE model (eqn. 4) is the only correct feeding model for our simulated data. Therefore, it is the only method which provides a jointly unbiased estimation of attack coefficients, handling times and attack exponents. As above, the methods of Bolker (eqn. 12) and Juliano (eqn. 11) underestimate attack coefficients due to the approximation of *a* = *bN*^*q*^ by 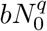. The deviation from all three true parameters is even stronger when directly fitting the functional response (eqn. 5) and Frost’s correction for prey depletion (eqn. 13) produces overall biased estimates.

### Prey mortality and prey growth

We compared three new ODE approaches of dealing with natural prey mortality (Fig. 3). When fitting a mortality-free model (eqn. 4) to data which includes natural mortality, attack coefficients and maximum feeding 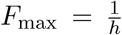 are systematically overestimated as compensation for the missing mortality term. The estimates of attack exponents are also biased as expected. Using a model including mortality (eqn. 14) leads to an unbiased prediction of all parameters. However, the error ranges for handling time *h* and mortality coefficient *m* are still relatively large, because effects of predation and natural mortality are correlated. But, if control data is used, the accuracy of predicted *h* and *m* is considerably improved, because predation and mortality effects can be disentangled.

**Figure 3:**
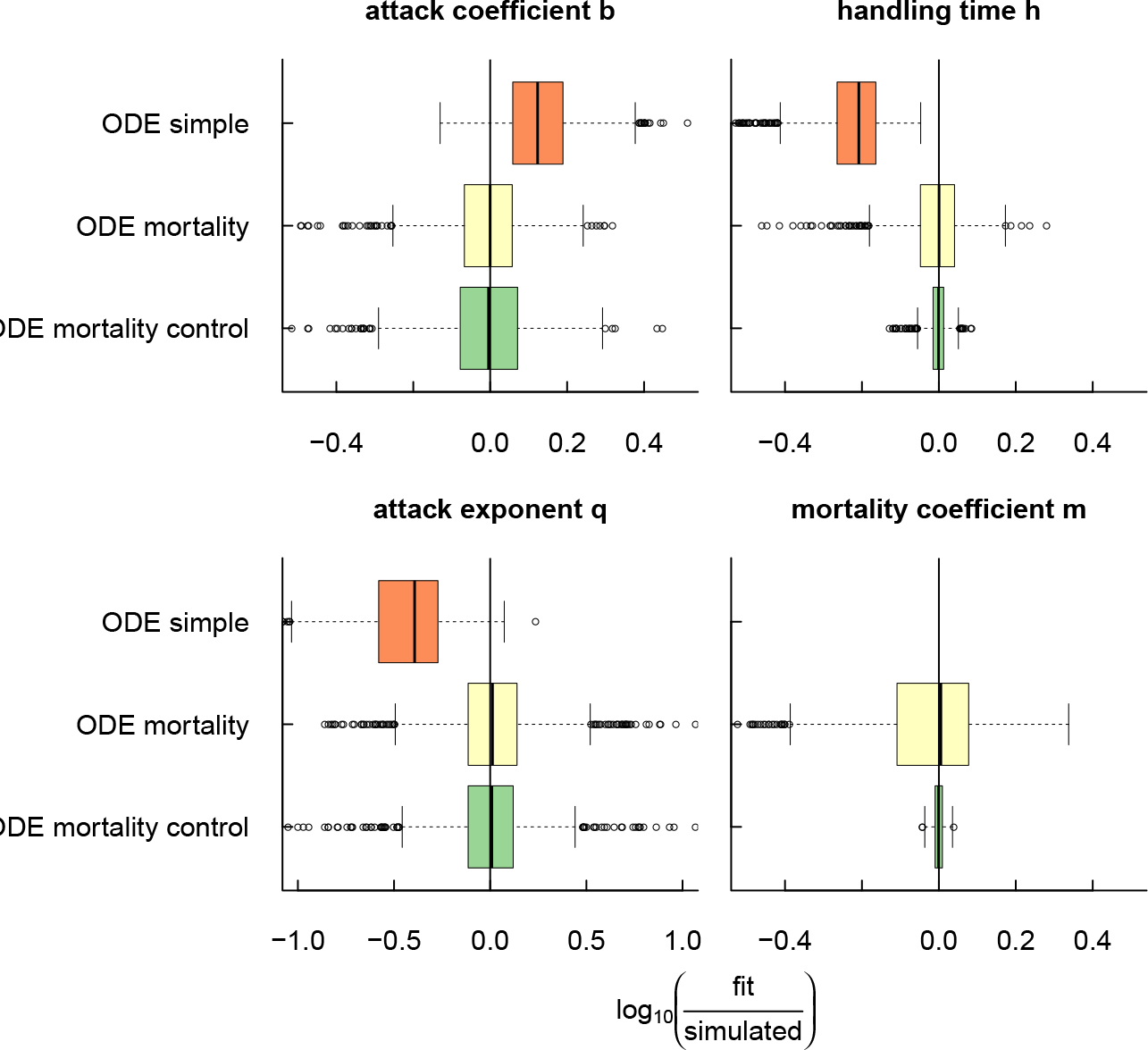
Evaluation of the ODE model with an additional prey mortality term. The boxplots show the error distributions of fitted parameters 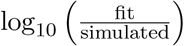 in 1000 simulated datasets.

Similarly, we compared three new ODE approaches of fitting data including natural prey growth and death (Fig. 4). Neglecting growth in the model (eqn. 4), handling times *h* are, unexpectedly, unbiased, but feature a comparatively large error range. Attack exponents *q* are, also unexpectedly, slightly underestimated. In combination with the severely underestimated attack coefficients *b* we assume that attack rates *a* = *bN*^*q*^ are generally underestimated as expected because they have to correct for the missing growth term in the model. When including this term (eqn. 15), all functional response parameters and also the growth term’s parameters, growth rate *r* and carrying capacity *K*, are unbiased. By fitting this growth model to data, which uses half of the number of observations as control data, the accuracy of the unbiased estimates is drastically improved.

**Figure 4:**
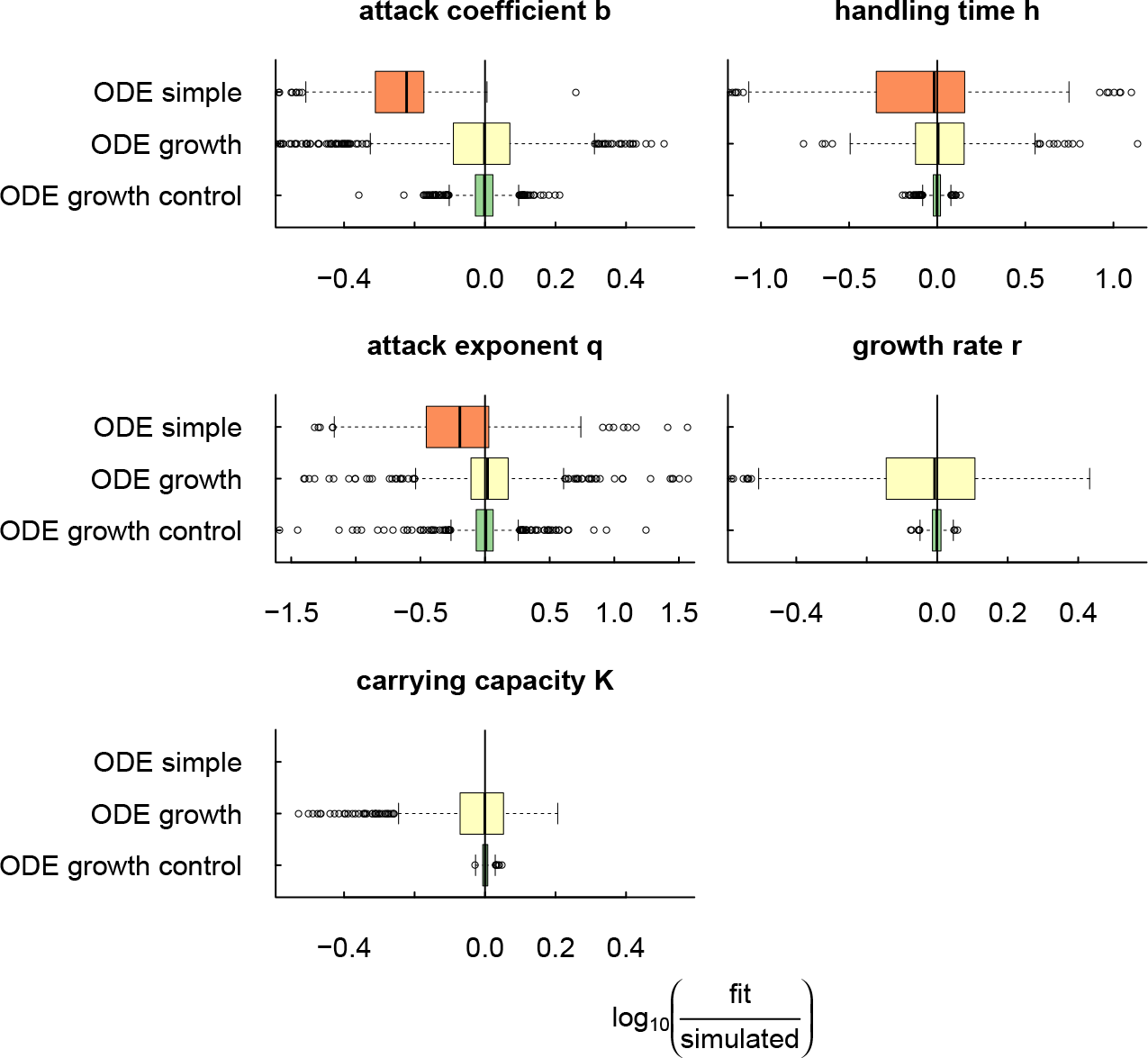
Evaluation of the ODE model with an additional prey growth and mortality term. The boxplots show the error distributions of fitted parameters 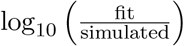 in 1000 simulated datasets.

### Experimental data

When fitting our new ODE model (eqn. 4) to experimental datasets, the predicted curves of eaten prey were almost identical to the studies’ original methods (Fig. 5). Both methods performed similarly in fitting the observed data. However, our focus lies on the parameters which produce these curves and on their discrepancies. Therefore, we do not provide model comparisons (e.g. AIC scores), but report these differences in the estimates (Table 1) and contrast them to our results found in simulated data.

**Figure 5:**
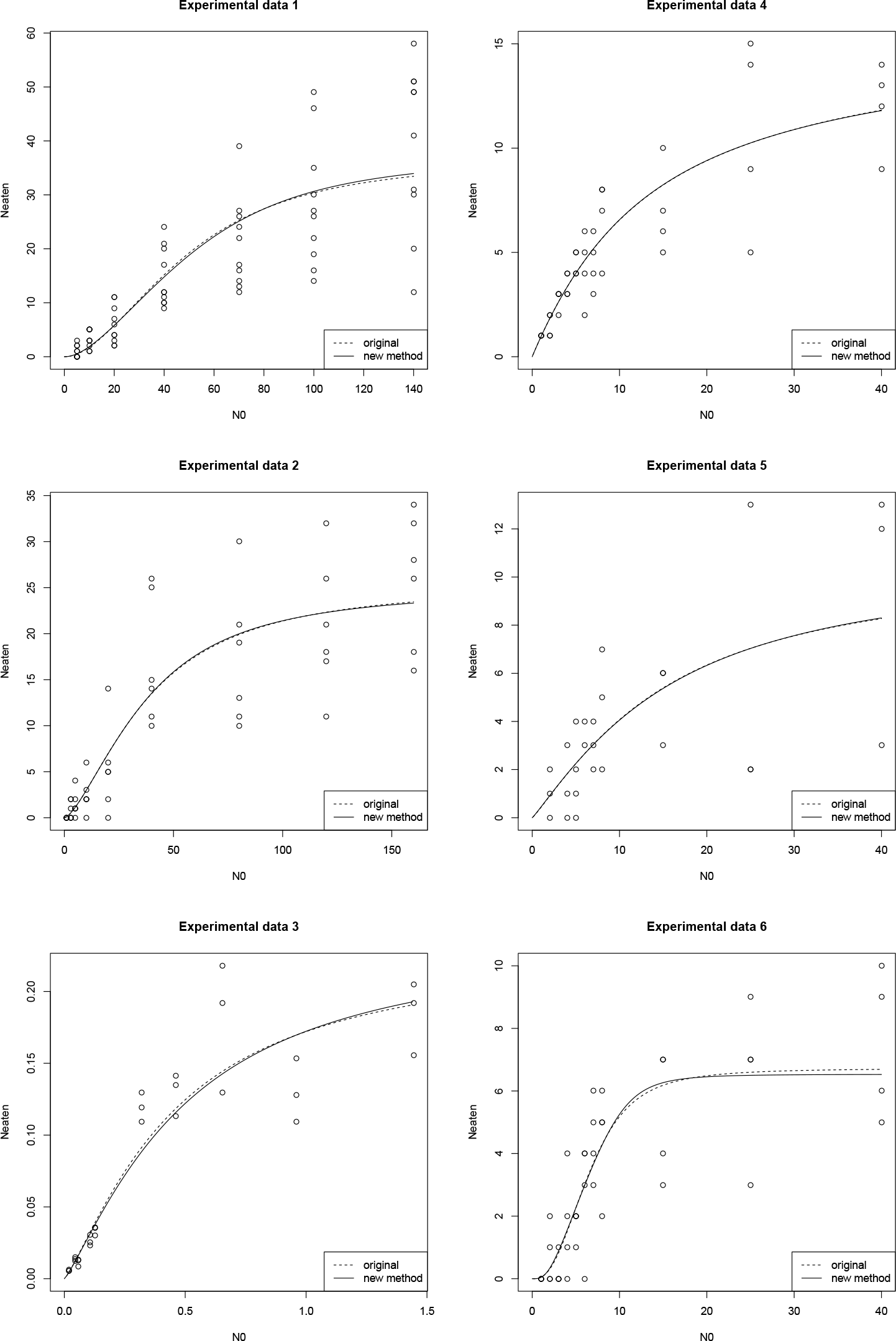
Fits of the original and our new ODE-based parameter estimation method to six experimental datasets D1-D6.

**Table 1:** Functional response parameters for experimental data (D1-D6). *n* = number of observations, *b* = attack coefficient, *h* = handling time, *q* = attack exponent. Standard errors are shown in parentheses. *P*-values: · *P* < 0.1, * P < 0.05, ** P < 0.01, *** P < 0.001.

| dataset | | method | $n$ | $b$ | $h$ | $q$ |
| --- | --- | --- | --- | --- | --- | --- |
| D1 (Sentis, Morisson & Boukal, 2015) | new | ODE III | 70 | 0.083 ***<br>($\pm 0.009$) | 0.008 ***<br>($\pm 0.000$) | |
| | old | RRPE III | 70 | 0.069 ***<br>($\pm 0.007$) | 0.008 ***<br>($\pm 0.000$) | |
| D2 (Vucic-Pestic *et al.*, 2010) | new | ODE gen | 54 | 0.112 *<br>($\pm 0.056$) | 0.040 ***<br>($\pm 0.004$) | 0.597 **<br>($\pm 0.197$) |
| | old | RRPE gen | 54 | 0.108 *<br>($\pm 0.052$) | 0.039 ***<br>($\pm 0.003$) | 0.565 **<br>($\pm 0.176$) |
| D3 (Uszko *et al.*, 2015) | new | ODE gen | 30 | 0.295 **<br>($\pm 0.095$) | 10.807 ***<br>($\pm 1.832$) | 0.247 *<br>($\pm 0.102$) |
| | old | Frost gen | 30 | 0.274 ***<br>($\pm 0.082$) | 10.426 ***<br>($\pm 1.897$) | 0.223 *<br>($\pm 0.095$) |
| D4 (Barrios-O’Neill *et al.*, 2015) | new | ODE gen | 44 | 2.688 ***<br>($\pm 0.350$) | 0.050<br>($\pm 0.032$) | -0.312<br>($\pm 0.236$) |
| | old | RRPE gen | 44 | 4.060 ·<br>($\pm 2.251$) | 0.051<br>($\pm 0.046$) | -0.410<br>($\pm 0.467$) |
| D5 (Barrios-O’Neill *et al.*, 2015) | new | ODE gen | 27 | 0.325<br>($\pm 0.230$) | 0.066 *<br>($\pm 0.027$) | 0.425<br>($\pm 0.461$) |
| | old | RRPE gen | 27 | 0.306<br>($\pm 0.227$) | 0.064 *<br>($\pm 0.026$) | 0.392<br>($\pm 0.415$) |
| D6 (Barrios-O’Neill *et al.*, 2015) | new | ODE gen | 44 | 0.019<br>($\pm 0.027$) | 0.172 ***<br>($\pm 0.021$) | 2.298 *<br>($\pm 0.961$) |
| | old | RRPE gen | 44 | 0.021<br>($\pm 0.024$) | 0.169 ***<br>($\pm 0.021$) | 1.937 **<br>($\pm 0.669$) |

The handling times *h* estimates are generally similar when comparing by the original and our new ODE method for all datasets. When dealing with a type III (dataset D1) or a generalized response (datasets D2, D3) with similarly fitted attack exponents *q*, the attack coefficients *b* are underestimated by the RRPE and also by Frost’s method, compared to the new ODE approach. This confirms our findings of the comparisons using simulated data and our assumptions about biased attack rates of the RRPE: By ignoring prey depletion and approximating *a* = *bN* by *bN*_0_ (or *a* = *bN*^*q*^ by 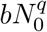 in the generalized case), *N*_0_ > *N* causes an underestimation of attack coefficients *b*. In datasets D4–D6, we observed both under-and overestimation of attack coefficients *b* by the RRPE-gen. Also, the attack exponents are underestimated. We emphasize that for generalized functional responses, attack coefficients and attack exponents in *a* = *bN*^*q*^ can only be interpreted jointly.

For the dataset D7 featuring natural prey mortality, we compared our generalized ODE model neglecting mortality (eqn. 4) to the ODE model including a mortality rate (eqn. 14) with and without using control data (Fig. 6, Table 2, see also the manual, Sec. 3). Again, the different models produce similar curves when fitting data with predators present, while the parameter estimates are quite different. As expected, the maximum feeding rate 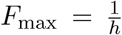 compensates for natural prey death and is therefore highly overestimated when fitting a model without mortality term, i.e. the handling time *h* is underestimated. When fitting the mortality model without using control data, a negative mortality rate *m* is predicted, which actually means natural prey growth instead of prey mortality. But the estimate is not significant and features a high standard deviation, because effects of feeding and natural prey mortality (or growth) cannot be disentangled. Using the same mortality model including additional control data, a positive mortality rate *m* is correctly estimated and we observe a reduced uncertainty (smaller standard error) in the estimates for *m*. Control data enables the precise separation of feeding and natural death.

**Figure 6:**
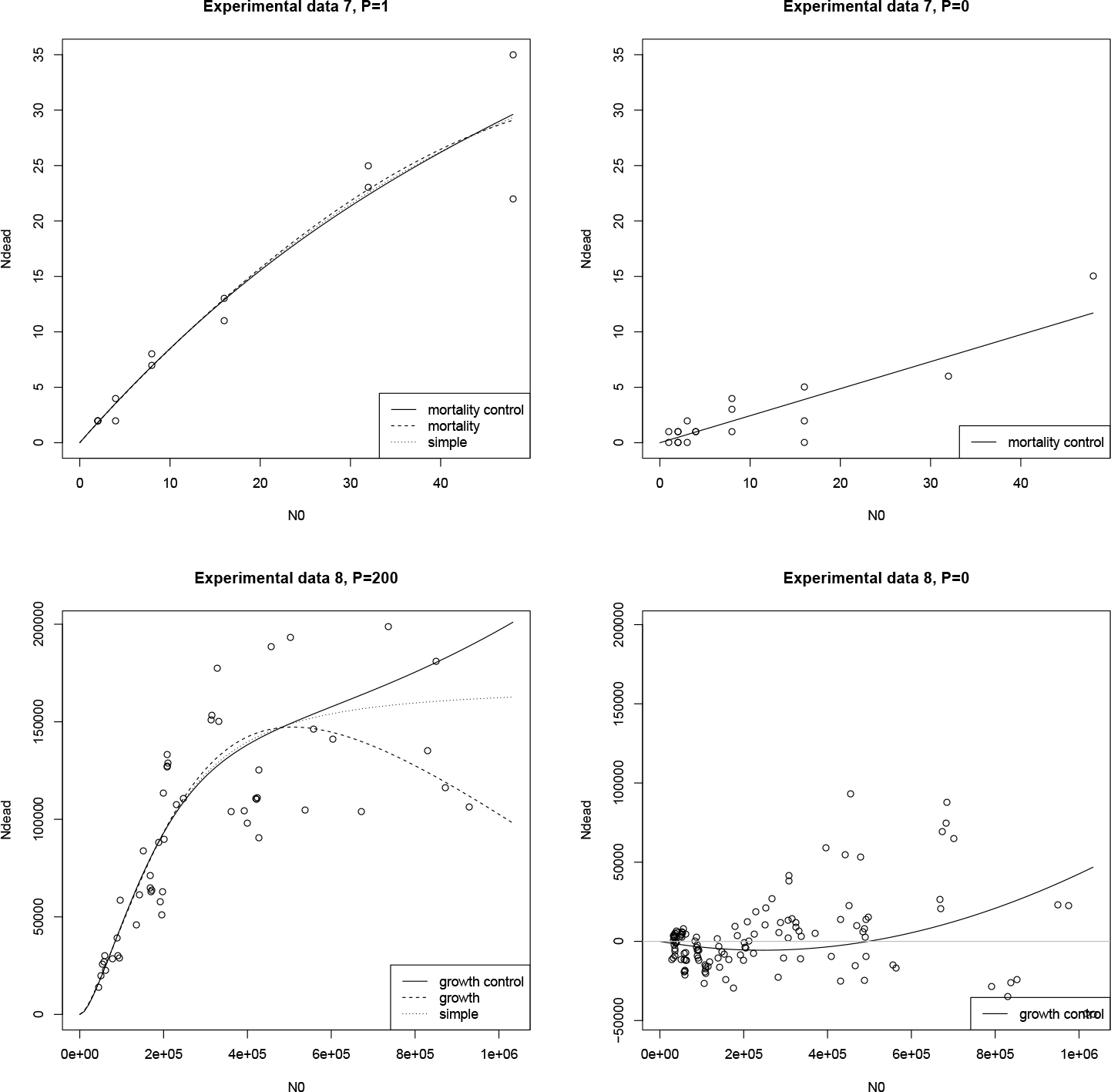
Fits of our new ODE-based models to experimental datasets D7 including background prey mortality (top) and D8 including background prey growth and mortality (bottom); feeding experiments (left) and control experiments (right).

**Table 2:** Functional response parameters for experimental data (D7, L. Archer, 2017, personal communication). *n* = number of observations, *b* = attack coefficient, *h* = handling time, *q* = attack exponent, *m* = mortality rate. Standard errors are shown in parentheses. P-values: · P < 0.1, * P < 0.05, ** P < 0.01, *** P < 0.001.

| | method | $n$ | $b$ | $h$ | $q$ | $m$ |
| --- | --- | --- | --- | --- | --- | --- |
| D7 | ODE | 13 | 2.415 ***<br>($\pm 0.699$) | 0.018<br>($\pm 0.014$) | -0.060<br>($\pm 0.272$) | |
| | ODE<br>+ mort | 13 | 3.329 *<br>($\pm 1.643$) | 0.005<br>($\pm 0.008$) | -0.062<br>($\pm 0.173$) | -0.995<br>($\pm 1.648$) |
| | ODE<br>+ mort + control | 31 | 2.207 **<br>($\pm 0.748$) | 0.031<br>($\pm 0.022$) | -0.049<br>($\pm 0.354$) | 0.279 ***<br>($\pm 0.043$) |

Dataset D8 includes natural prey growth and death and we compared our generalized ODE model ignoring background growth (eqn. 4) to the ODE model including a logistic growth term (eqn. 15) with and without using control data (Fig. 6, Table 3). Note that the parameters *r*, *K*, *b* and *h* are estimated on their log scale. If parameters differ by several orders of magnitude, this can improve the efficiency of the iterative maximum likelihood algorithm; see the manual (Secs. 5, 6.6) for details. Surprisingly, the parameter estimates for fitting the model neglecting growth and the full model including control data are similar, although their predictions do differ for large initial prey densities. Fitting the growth model without control data produces heavily-biased estimates. A large carrying capacity *K* and a large growth rate *r* lead to a decrease in predicted dead prey for large initial prey densities. Using control data, the uncertainty in all parameters is significantly reduced (smaller standard errors), because effects of feeding and natural growth (*N* < *K*) (or natural loss, *N* > *K*) can be separated precisely.

**Table 3:** Functional response parameters for experimental data (D8, Fussmann, 2017). *n* = number of observations, *b* = attack coefficient, *h* = handling time, *q* = attack exponent, *r* = growth rate, *K* = carrying capacity. Standard errors are shown in parentheses. P-values: · P < 0.1, * P < 0.05, ** P < 0.01, *** P < 0.001.

| | method | $n$ | $\log(b)$ | $\log(h)$ | $q$ | $\log(r)$ | $\log(K)$ |
| --- | --- | --- | --- | --- | --- | --- | --- |
| D8 | ODE | 53 | -19.356 ***<br>($\pm 2.954$) | -1.250 ***<br>($\pm 0.146$) | 0.751 **<br>($\pm 0.267$) | | |
| | ODE<br>+ growth | 53 | -16.387 ***<br>($\pm 3.217$) | -1.768 ***<br>($\pm 0.535$) | 0.499 ·<br>($\pm 0.283$) | -7.238 ***<br>($\pm 0.485$) | 16.934<br>NA |
| | ODE<br>+ growth + control | 179 | -19.401 ***<br>($\pm 2.181$) | -1.257 ***<br>($\pm 0.111$) | 0.763 ***<br>($\pm 0.198$) | -8.580 ***<br>($\pm 0.376$) | 13.115 ***<br>($\pm 0.346$) |

## Discussion

We compared different popular methods and our new ODE-based approach for fitting functional response models to feeding experiments, using both simulated and experimental datasets. We estimated parameters of type II, type III and the generalized functional response models via maximum likelihood. Additionally, we investigated models including background prey mortality or growth. Based on our results, we give the following recommendations:

While our new ODE-approach generally yields unbiased estimates by correcting for prey depletion, direct methods (without requiring numerical simulation) perform equally when using the type II (*q* = 0) or the strict type III model (*q* = 1). Bolker’s explicit solution (Bolker, 2008) of the RRPE-II (eqn. 7) can be used for fitting type II responses. For type III responses, we propose using our explicit version (eqn. 10) of Hassell’s approach (Hassell, Lawton & Beddington, 1977) as the fit of an analytical solution is way faster than fitting simulations to data (see supplement, Sec. 6).

However, when including the attack exponent *q* as a free parameter, only numerical simulation of ODEs guarantees unbiased estimates. This is of utmost importance for ecological research as even a small shift of the attack exponent, *q*, from 0 (strict type II functional response) to approximately 0.2 (e.g. Williams & Martinez, 2004) increases stability of trophic ecological networks (i.e. food webs) dramatically and thereby also increases species coexistence and biodiversity (Rall, Guill & Brose, 2008). Studies investigating empirically the attack exponent increased in number over the last years (Vucic-Pestic *et al.*, 2010; Kalinkat *et al.*, 2013; Barrios-O’Neill *et al.*, 2015, 2016; Uszko *et al.*, 2015, 2017). Such publications will benefit from our new approach as it allows ecologists to correctly estimate the functional response parameters that are needed to gain the understanding of stability and diversity patterns in complex ecological communities.

Besides these obvious and needed advancements, our ODE-based method is highly flexible. It is not limited to the three functional response models described here - any type of functional response describing the prey-dependent feeding rate can be integrated. Using simulations of ODEs that are fitted to data further allows ecological modelers to include additional terms like prey mortality or growth, as presented in this study. We showed, however, that control data is required to disentangle effects of natural mortality or growth and predation.

Additionally to this main article, we provided a in-depth description of our method as a manual, R-source files, and data. This will allow researchers that want to apply our method to their own functional response data an easy entry into our methodology and the possibility to practice using the method with worked examples before using it to analyze their own data. Moreover, the source files attached in the supplement can be adapted to include other functional response models than those used here (see Juliano (2001) and Jeschke, Kopp & Tollrian (2002) for a variety of possible models). In addition, specific mortality and growth models can be incorporated, e.g. using the Gompertz growth model instead of the logistic growth model - a growth model often used in microbiology (Paine *et al.*, 2012; Fussmann *et al.*, 2017).

We also show in the supplement (Sec. 7) how data availability changes the outcome of the parameter fitting. Generally, we found that even sparse data is sufficient to estimate the parameters correctly, but especially the estimation of handling time is highly uncertain if higher densities of prey are not measured in an experiment (Supp. Fig. 2).

In conclusion, we found that our new method to estimate functional response parameters from a feeding experiment with prey depletion - the decrease of prey number over time without replacement - is superior to traditional approaches. To our knowledge, it is the only method that allows for precise parameter estimation of the generalized functional response, which is currently one of the most frequently-used descriptions of feeding interactions in ecology. It moreover allows easy adaptation and inclusion of unwanted effects like prey growth and mortality, what was impossible with traditional approaches. We believe that our approach to estimate functional response parameters will enable researchers in the future to estimate precise parameter values, using mortality-and growth-biased data, and thereby deepen understanding of how interactions drive stability and biodiversity of complex ecological communities.

## Acknowledgements

We gratefully acknowledge the support of the German Centre for integrative Biodiversity Research (iDiv) Halle-Jena-Leipzig funded by the German Research Foundation (FZT 118). We thank Christian Guill for providing the analytical derivation of the RRPE from an ODE including a type II functional response. We thank Arnaud Sentis as well as Louise Archer, Esra Sohlström, Bruno Gallo, Malte Jochum, Guy Woodward, Rebecca L. Kordas, Eoin J. O’Gorman and Katarina Fussmann for providing data that we could re-analyse. We are grateful for authors that make their data freely avaible from Barrios-O’Neill *et al.* (2015); Uszko *et al.* (2015); this allowed us to perform this study. Last but not least we want to thank Callum Lawson and Gregor Kalinkat for proof reading and helpful comments on our manuscript.

## Data accessibility

We attached the empirical data used in that study as supplemental information together with an in-depth description of the fitting methods in the manual, including R-source files allowing to reproduce our method.

